# TCA Cycle Dysfunction and Amino Acid Catabolism Drive Hepatic Steatosis in Mice with HFpEF

**DOI:** 10.1101/2024.06.03.597212

**Authors:** Bellina A.S. Mushala, Michael W. Stoner, Janet R. Manning, Paramesha Bugga, Nisha Bhattarai, Maryam Sharifi-Sanjani, Brenda McMahon, Amber Vandevender, Steven J. Mullet, Brett A. Kaufman, Sruti S. Shiva, Stacy L. Gelhaus, Michael J. Jurczak, Iain Scott

## Abstract

The prevalence of cardiometabolic heart failure with preserved ejection fraction (HFpEF) continues to grow worldwide, and now represents over half of current heart failure cases in the United States (1). Due to a lack of specific approved therapies, current treatment guidelines focus on the management of comorbidities related to metabolic syndrome (e.g. obesity, diabetes, hypertension) that promote HFpEF progression (1). The same comorbidities also drive cardiometabolic disease in non-cardiac tissues, and links between disease presentations in different organs are increasingly being recognized in the clinic. However, mechanistic studies examining the underlying pathophysiological connections have not kept pace, particularly in the cardio-hepatic disease axis (2). To address this, we used a recently developed and validated preclinical model of HFpEF (3) to examine how this disease impacts the liver. The development of HFpEF in mice leads to the simultaneous development of widespread hepatic steatosis that is consistent with human non-alcoholic fatty liver disease (NAFLD). Mechanistically, we show that the liver steatosis observed is driven by excess glucogenic amino acid entry into the TCA cycle, which promotes hepatic glucose production and de novo lipogenesis. Our findings suggest that HFpEF development is a multi-organ event, with implications for both preclinical and translational research.

## Results and Discussion

Adult male C57BL/6J mice were fed either a control chow diet, or a combination of 60% high fat diet plus 0.5 g/L L-NAME in drinking water for 15 weeks to induce HFpEF (**Figure 1A**). HFpEF mice displayed increased body weight, fat mass, and insulin resistance (**Figure 1B-D**). Heart weight was elevated in HFpEF mice (**Figure 1E**), and as previously reported this model induced diastolic dysfunction in the absence of systolic impairment (**Figure 1F,G**). Histology showed that HFpEF mice had significantly elevated levels of hepatic steatosis and inflammation (**Figure 1 H-J**), which were consistent with non-alcoholic fatty liver disease (NAFLD) in humans. Regression analyses demonstrated that histological NAFLD activity scores (NAS) correlated with diastolic, but not systolic, cardiac dysfunction (**Figure 1K**). Bulk liver transcriptomic analysis showed differentially expressed genes (DEGs; **Figure 1L; Table S1**) that were predominantly linked to cytochrome P450, fatty acid, and metabolite transport processes (**Figure 1M-P**). Despite the significant increase in liver steatosis, there were no overall changes in the expression or abundance of key liver fatty acid uptake proteins or oxidation enzymes (**Figure 1O, Figure S1**). Notably, there was little overlap of DEGs identified here and previous transcript profiling of human NAFLD (**Figure S2**) (4), which suggests that a HFpEF-specific expression profile may exist.

**Figure 1:**
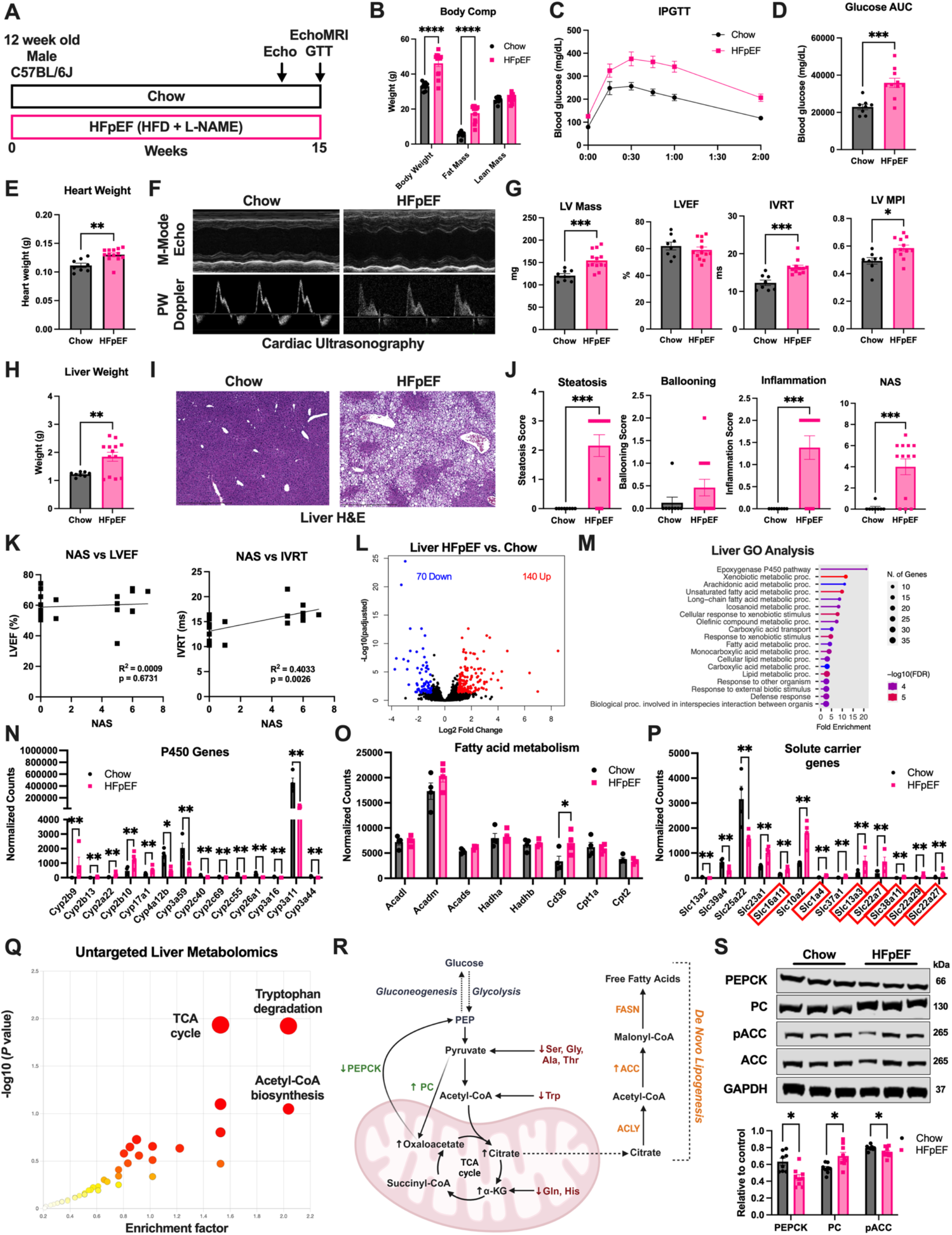
Hepatic TCA cycle dysfunction drives steatosis. (A) Schematic of mouse HFpEF model. (B) EchoMRI measurements of total body weight, fat mass, and lean mass. (C,D) Intraperitoneal glucose tolerance test (IPGTT) and glucose area under the curve (AUC). (E) Heart weight. (F) M-mode echocardiography and pulsed-wave (PW) Doppler of cardiac function. (G) Echocardiographic measurement of left ventricle (LV) mass, ejection fraction (LVEF), isovolumetric relaxation time (IVRT), and myocardial performance index (LV MPI). (H) Liver weight. (I) H&E histology images of liver. (J) Histological scoring of steatosis, ballooning, inflammation, and total NAFLD activity score (NAS). (K) Regression analysis of NAS and either LVEF or IVRT. (L) Transcriptomic analysis of chow and HFpEF livers. Genes with an adjusted *P* < 0.05 and absolute log2 FC > 1 were designated as differentially expressed genes (DEGs). (M) Gene ontology analysis of liver DEGs. (N) Relative expression of Cytochrome P450 genes. (O) Relative expression of fatty acid metabolism genes. (P) Relative expression of solute carrier genes. Genes involved in amino acid or carboxylate transport are highlighted in red. (Q) Untargeted metabolomics analysis of liver tissue, with top three enriched pathways annotated. (R) Summary of alterations in TCA cycle, cataplerosis, and de novo lipogenesis in livers from HFpEF mice. (S) Western blot analysis of phosphoenolpyruvate carboxykinase (PEPCK), pyruvate carboxylase (PC), and phosphorylated acetyl-CoA carboxylase (pACC). Data presented as mean ± SEM. Unpaired student’s t-test was used for comparisons between two groups. * = *P* < 0.05; ** = *P* < 0.01; *** = *P* < 0.001; **** = *P* < 0.0001.

We next performed untargeted metabolomics to understand what pathways may be contributing to the observed liver steatosis, and found that TCA cycle processes, amino acid degradation, and acetyl-CoA biosynthesis were most enriched (**Figure 1Q; Table S2**). To dissect these changes, we performed targeted metabolomics on bioenergetic substrates and TCA cycle intermediates. Glucose and glycolysis intermediates were unchanged between chow and HFpEF animals (**Figure 1R, Figure S1**). In contrast, there was a significant increase in several TCA cycle intermediates in HFpEF livers (oxaloacetate, citrate, cis-aconitate, and α-ketoglutarate), while others (succinate, fumarate, malate) remained unchanged (**Figure 1R, Figure S1**). Strikingly, increased TCA cycle intermediates correlated with a significant decrease in glucogenic amino acids that enter the glycolysis/TCA cycle pathways at pyruvate (serine, glycine, alanine, and threonine), acetyl-CoA (tryptophan), or α-ketoglutarate (glutamine, histidine) (**Figure 1R, Figure S1**). These changes correlated with an increase in solute carrier genes involved in amino acid or carboxylate transport (**Figure 1P**), which combined suggests an increase in amino acid catabolism in HFpEF livers.

Liver pyruvate carboxylase (PC) was elevated in HFpEF mice which, combined with elevated fasting blood glucose levels, suggested increased hepatic glucose production (**Figure 1C,R,S**). However, a corresponding decrease in phosphoenolpyruvate carboxykinase (PEPCK) likely prevented the complete diversion of TCA intermediates into the gluconeogenesis pathway (**Figure 1S**). Instead, elevated TCA cycle intermediates appeared to contribute to *de novo* lipogenesis, as demonstrated by a significant reduction in the inhibitory phosphorylation of acetyl-CoA carboxylase (ACC) (**Figure 1R,S**). Overall, we conclude that TCA cycle dysfunction driven by elevated amino acid catabolism in the livers of HFpEF mice promotes gluconeogenesis and hepatic steatosis.

There are two main caveats to our study. Firstly, female sex has been shown to be protective in this preclinical HFpEF model (5), therefore our findings may be specific to males analyzed here. Secondly, our metabolomics study was cross-sectional, therefore flux measurements were not possible. While TCA cycle dysfunction has been noted previously in human NAFLD (6), this is the first report of increased amino acid catabolism fueling both gluconeogenesis and *de novo* hepatic lipogenesis in HFpEF. Combined, our findings highlight the need for the development of HFpEF treatments that take account of the importance of multi-organ disease.

