## Supplemental Figures for "TCA Cycle Dysfunction and Amino Acid Catabolism Drive Hepatic Steatosis in Mice with HFpEF"

### SUPPLEMENTAL METHODS

#### Animal Care and Use

Male C57BL/6J 8-week-old mice were obtained from the Jackson Laboratory and maintained on chow for 4 weeks to acclimate to their new environment. Animals were housed in the University of Pittsburgh animal facility under standard conditions, with ad libitum access to water and food, and maintained on a constant 12-hour light/dark cycle. At 12 weeks of age, mice were exposed to normal chow (Chow; 60% carbohydrate, 26% protein, 14% fat; ProLab IsoPro RMH 3000), or a combination of high-fat diet (HFD; 20% carbohydrate, 20% protein, 60% fat; Research Diets D12492) plus Nω-nitro-L-arginine methyl ester (L-NAME: 0.5 g/L, pH 7.4; ThermoFisher) supplemented in their drinking water to induce HFpEF (Schiattarella et al., 2019). Mice were subject to the HFpEF diet for 15 weeks. Animals were euthanized by isofluorane anesthesia followed by cervical dislocation. Experiments were conducted in compliance with National Institutes of Health guidelines, and followed procedures approved by the University of Pittsburgh Institutional Animal Care and Use Committee.

#### Echocardiography

Mice were anesthetized using isofluorane (1.5%–2.0% v/v by inhalation) and monitored for cardiac functional parameters in the supine position using a Visual Sonics Vevo 3100. Core temperature was maintained at 37 °C by imaging mice while on a heating pad, and heart rates were kept consistent between experimental groups (~400–500 beats per min). Short axis M-mode was used to assess LV dimensions and motion patterns, pulsed-wave (PW) doppler mode to evaluate blood flow velocities across the mitral valve, and tissue doppler mode to measure mitral annular plane velocity. Doppler profiles were acquired in the parasternal long axis (PLAX) apical 4 chamber view. The left atrial area was quantified in the apical 4-chamber view by tracing the border of the left atrium. Markers of systolic and diastolic function were calculated using standard echocardiography equations. At the end of the procedure, all mice recovered from anesthesia without difficulties. Image analysis was performed independently by a blinded sonographer and all parameters were measured at least 3 times, and averages are presented.

**Histology**

Preparation and staining of all histological samples were conducted by the Pitt Biospecimen Core at the University of Pittsburgh. All mice were sacrificed, and tissue harvested for analysis within one hour of completion of the IPGTT study. The left lobe of the liver was excised from each mouse following euthanasia, fixed in 10% buffered formalin phosphate overnight at shaking incubation, washed 3 times for 5 min with 1X phosphate-buffered saline (PBS), and then transferred to 70% ethanol. Samples were then embedded in paraffin, sectioned into 4 µm slides, and stained with hematoxylin and eosin (H&E). The NAFLD activity score (NAS) was obtained based on methods established by the Pathology Committee of the Non-alcoholic Steatohepatitis (NASH) Clinical Research Network (Kleiner et al., 2005). Liver pathology scoring was performed in a blinded independent fashion. NAS consisted of separate category scores which included steatosis (0-3), where 0 = 0–5%, 1 = 6–33%, 2 = 34–66%, and 3 = 67–100% of hepatocytes positive for steatosis; lobular inflammation (0-3), where 0 = no foci, 1 = more than 2 foci, 2 = 2-4 foci and 3 = greater than 4 foci all per 200X field; and hepatocellular ballooning (0-2), where 0 = none present, 1 = few and 2 = many/prominent. The final NAS represents a sum of the three category scores. Histology sections were visualized with the Evos FL Auto 2 Microscope, observed changes in structure were quantified using ImageJ software and representative images are shown.

**Transcriptomics**

Total RNA was isolated from hepatic tissue using the RNeasy Plus Mini Kit (Qiagen REF 74134). Approximately 2 µg RNA was used for bulk RNA sequencing performed by Azenta/GENEWIZ, based on company recommendations to identify differential gene expression patterns between the experimental groups.

**Quantitative Metabolomics**

Metabolic quenching and polar metabolite pool extraction was performed by adding ice cold 80% methanol (aqueous) at a ratio of 1:15 wt input tissue:vol. ($^{13}C_1$)-creatinine, ($D_3$)-taurine, ($D_3$)-lactate and ($D_3$)-alanine (Sigma-Aldrich) were added to the sample lysates as an internal standard for a final concentration of 10 µM. Samples are homogenized using an MP Bio FastPrep system using Matrix D (ceramic sphere) for 60 s at 60 hz. The supernatant was then cleared of protein by centrifugation at 16,000 *g*. 2 µL of cleared supernatant was subjected to online LC-MS analysis. Analyses were performed by untargeted liquid chromatography-high-resolution mass spectrometry (LC-HRMS). Briefly, samples were injected via a Thermo Vanquish UHPLC and separated over a reversed phase. Thermo HyperCarb porous graphite column (2.1×100 mm, 3 µm particle size) maintained at 55 °C. For the 20 min LC gradient, the mobile phase consisted of the following: solvent A (water/0.1% FA) and solvent B (ACN/0.1% FA). The gradient was the following: 0-1 min 1% B, increase to 15% B over 5 min, continue increasing to 98% B over 5 min, hold at 98% B for 5 min, re-equillibrate at 1% B for 5 min. The Thermo IDX tribrid mass spectrometer was operated in both positive and negative ion mode, scanning in ddMS2 mode (2 µscans) from 70 to 800 m/z at 120,000 resolution with an AGC target of $2e^5$ for full scan, $2e^4$ for $ms^2$ scans using HCD fragmentation at stepped 15, 35, 50 collision energies. Source ionization setting was 3.0 and 2.4 kV spray voltage, respectively, for positive and negative mode. Source gas parameters were 35 sheath gas, 12 auxiliary gas at 320 °C, and 8 sweep gas. Calibration was performed prior to analysis using the PierceTM FlexMix Ion Calibration Solutions (Thermo Fisher Scientific). Integrated peak areas were then extracted manually using Quan Browser (Thermo Fisher Xcalibur ver. 2.7). Untargeted differential comparisons were performed using Compound Discoverer 3.0 (Thermo Fisher) to generate a ranked list of significant compounds with tentative identifications from BioCyc, KEGG, and internal compound databases. Purified standards were then purchased and compared in retention time, m/z, along with $ms^2$ fragmentation patterns to validate the identity of significant hits.

**Protein Isolation and Immunoblotting**

Liver tissues were rapidly harvested following euthanasia, weighed, and flash-frozen in liquid nitrogen. For protein isolation, tissues were minced and lysed in CHAPS buffer (1% CHAPS, 150 mM NaCl, 10 mM HEPES, pH 7.4) using a VWR 4-Place Mini Bead Mill, then incubated on ice for ~ 2.5 h. Homogenates were spun at 10,000 *g* at 4 °C for 10 min, and the supernatants collected for immunoblotting. For immunoblotting, protein lysates were quantitated using a BioDrop µLITE Analyzer, prepared in LDS sample buffer, separated using Bolt SDS-PAGE 4–12% or 12% Bis–Tris Plus gels, and transferred to nitrocellulose membranes (all Invitrogen). Membranes were blocked using SuperBlock (PBS) Blocking Buffer and incubated overnight in the following primary antibodies: rabbit phos-ACC (Ser79) (11818S), rabbit ACC (3676P), rabbit phos-ACLY (Ser455) (4331S), rabbit ACLY (13390S) and rabbit GAPDH (2118S) from Cell Signaling; rabbit DGAT2 (ab96094) from Abcam; rabbit PDK4 (PA5-13776) from ThermoFisher; rabbit CD36 (18836-1-AP), rabbit CPT1a (15184-1-AP), rabbit HADHA (10758-1-AP), rabbit LCAD (17526-1-AP), rabbit PEPCK1 (16754-1-AP), rabbit PC (16588-1-AP) and rabbit G6PC (22169-1-AP) from ProteinTech. Protein loading was confirmed using GAPDH as a loading control. Fluorescent anti-goat or anti-rabbit secondary antibodies (red, 700 nm; green, 800 nm) from LiCor were used to detect expression levels. Images were obtained using Licor Odyssey CLx System and protein densitometry was measured using the LiCor Image Studio Lite Ver. 5.2 Software.

**Statistics**

Means ± SEM were calculated for all data sets. Data comparisons between two groups were analyzed using the Student's t-test. Time course data was analyzed using two-way ANOVA with Sidak's post-hoc testing. $P \leq 0.05$ was considered statistically significant. Statistical analyses were performed using GraphPad Prism 9.5 Software.

SUPPLEMENTAL FIGURES

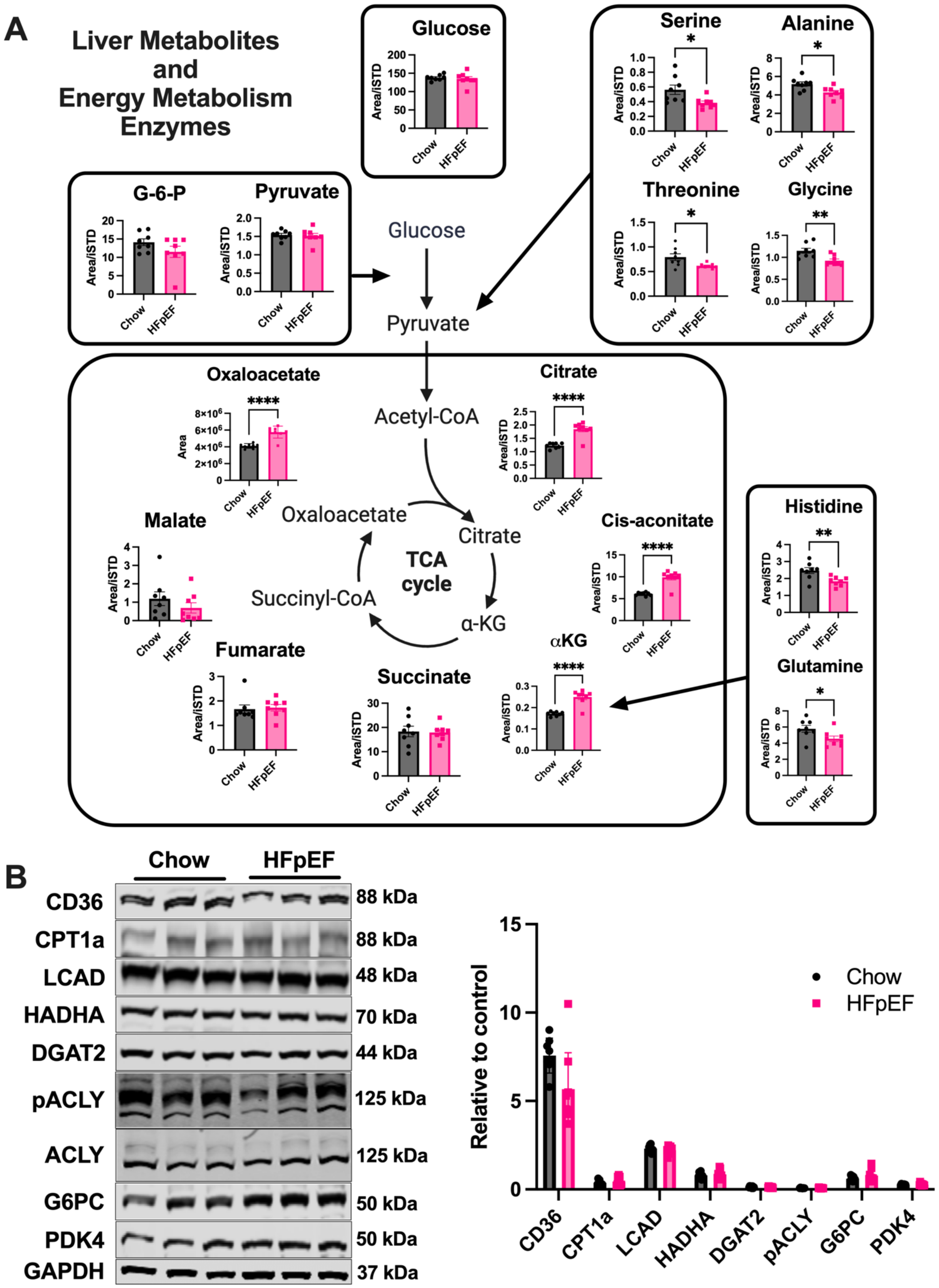

**Supplemental Figure 1: Energy metabolism analysis in livers from chow and HFpEF mice.** (A) Untargeted (oxaloacetate) and targeted (all other metabolites) metabolomics measurements from liver tissue excised from chow and HFpEF mice. (B) Western blot analysis of fatty acid and glucose metabolism proteins from chow and HFpEF livers. Unpaired student's t-test was used for comparisons between two groups. * = $P < 0.05$; ** = $P < 0.01$; *** = $P < 0.001$; **** = $P < 0.0001$.

| Gene | Fold change | p value (adj) |
|---|---|---|
| Akr1b10 | 0.006985368 | 0.998855265 |
| Ankrd29 | nd | |
| Ccl20 | nd | |
| Cfap221 | nd | |
| Clic6 | nd | |
| Col1a1 | 0.74177072 | 0.407305314 |
| Col1a2 | 0.733497159 | 0.128603155 |
| Dtna | nd | |
| Dusp8 | 0.535545067 | 0.631400575 |
| Epb41l4a | -0.299325469 | 0.86868388 |
| Fermt1 | -0.477164344 | 0.900446866 |
| Gdf15 | 1.701556828 | 2.13E-06 |
| Hecw1 | nd | |
| Hsd17b14 | nd | |
| Il32 | nd | |
| Itgbl1 | nd | |
| Ltbp2 | nd | |
| Pdgfa | 0.171537664 | 0.910165402 |
| Ppapdc1a | nd | |
| Rgs4 | 1.264166337 | 0.138998851 |
| Sctr | nd | |
| Stmn2 | nd | |
| Thy1 | -0.001223232 | 0.999377524 |
| Tnfrsf12a | -0.20138953 | 0.912297734 |
| Tyms | 0.023431884 | 0.996124051 |

**Supplemental Figure 2: Comparison of mouse HFpEF liver genes with human NAFLD gene signature.** *Gdf15* is the only transcript to be significantly elevated in both mouse HFpEF livers, and in human NAFLD reported by Govaere et al. (3). nd = no data.

**ACKNOWLEDGEMENTS**

This work was supported by: National Institute of Health F31 Fellowship (1F31DK134089-01A1) to B.A.S.M; National Institute of Health Shared Instrumentation Grant (S10OD023402, S10OD032141) to S.L.G., and National Institute of Health research grant (R01HL147861, R0HL156874) to I.S. The University of Pittsburgh Center for Metabolism and Mitochondrial Medicine is supported by the Pittsburgh Foundation (MR2020 109502) grant to M.J.J. This project used the University of Pittsburgh Medical Center (UPMC) Hillman Cancer Center and Tissue and Research Pathology/Pitt Biospecimen Core shared resource, which is supported in part by award P30CA047904. Echocardiography was carried out by the University of Pittsburgh Rodent Ultrasonography Core, which received funding from the NIH Shared Instrumentation Grant Program (1S10OD023684-01A1; Advanced High Resolution Rodent Ultrasound Imaging System).

### AUTHOR CONTRIBUTIONS

B.A.S.M., M.J.J., and I.S. conceived the study and designed experiments. B.A.S.M, M.W.S., B.M., A.M.V., and S.J.M. performed experiments. B.A.S.M., B.M., S.J.M. and I.S. analyzed data. B.A.S.M. and I.S. prepared figures. J.R.M., P.B., N.B., M.S.S., S.S.S., B.A.K., and S.L.G provided critical input and expertise. B.A.S.M. and I.S. drafted the manuscript. B.A.S.M., M.J.J., and I.S. edited and revised the manuscript.
